# Clomipramine suppresses ACE2-mediated SARS-CoV-2 entry

**DOI:** 10.1101/2021.03.13.435221

**Authors:** Yuri Kato, Shigeru Yamada, Kazuhiro Nishiyama, Ayano Satsuka, Suyong Re, Daiki Tomokiyo, Jae Man Lee, Tomohiro Tanaka, Akiyuki Nishimura, Kenzo Yonemitsu, Hiroshi Asakura, Yuko Ibuki, Yumiko Imai, Noriho Kamiya, Kenji Mizuguchi, Takahiro Kusakabe, Yasunari Kanda, Motohiro Nishida

**Affiliations:** Graduate School of Pharmaceutical Sciences, Kyushu University, Fukuoka 812-8582, Japan; Division of Pharmacology, National Institute of Health Sciences (NIHS), Kanagawa, 210-9501, Japan; Artificial Intelligence Center for Health and Biomedical Research (ArCHER), National Institutes of Biomedical Innovation, Health and Nutrition, 7-6-8 Saito-Asagi, Ibaraki, Osaka, 567-0085, Japan; Laboratory of Creative Science for Insect Industries, Faculty of Agriculture, Kyushu University, Motooka 744, Nishi-ku, Fukuoka, 819-0395, Japan; National Institute for Physiological Sciences & Exploratory Research Center on Life and Living Systems, National Institutes of Natural Sciences, Okazaki, 444-8787, Japan; Division of Biomedical Food Research, National Institute of Health Sciences (NIHS), Kanagawa, 210-9501, Japan; Graduate Division of Nutritional and Environmental Sciences, University of Shizuoka, 52-1 Yada, Shizuoka 422-8526, Japan; Laboratory of Regulation for Intractable Infectious Diseases, Center for Vaccine and Adjuvant Research (CVAR), National Institutes of Biomedical Innovation Health and Nutrition (NIBIOHN), Osaka, 567-0085, Japan; Department of Applied Chemistry, Graduate School of Engineering, Kyushu University, Fukuoka, 819-0395, Japan; Division of Biotechnology, Center for Future Chemistry, Kyushu University, Fukuoka, 819-0395, Japan; Institute for Protein Research, Osaka University, 3-2 Yamadaoka, Suita, Osaka, 565-0871, Japan; Laboratory of Insect Genome Science, Faculty of Agriculture, Kyushu University, Motooka 744, Nishi-ku, Fukuoka, 819-0395, Japan

**Author notes:** Correspondences (Y.K.), (M.N.).

## Abstract

Myocardial damage caused by the newly emerged coronavirus (SARS-CoV-2) infection is one of key determinants of COVID-19 severity and mortality. SARS-CoV-2 entry to host cells are initiated by binding with its receptor, angiotensin converting enzyme (ACE) 2, and the ACE2 abundance is thought to reflect the susceptibility to infection. Here, we found that clomipramine, a tricyclic antidepressant, potently inhibits SARS-CoV-2 infection and metabolic disorder in human iPS-derived cardiomyocytes. Among 13 approved drugs that we have previously identified as potential inhibitor of doxorubicin-induced cardiotoxicity, clomipramine showed the best potency to inhibit SARS-CoV-2 spike glycoprotein pseudovirus-stimulated ACE2 internalization. Indeed, SARS-CoV-2 infection to human iPS-derived cardiomyocytes (iPS-CMs) and TMPRSS2-expressing VeroE6 cells were dramatically suppressed even after treatment with clomipramine. Furthermore, the combined use of clomipramine and remdesivir was revealed to synergistically suppress SARS-CoV-2 infection. Our results will provide the potentiality of clomipramine for the breakthrough treatment of severe COVID-19.

## Introduction

A new pandemic of pneumonia caused by severe acute respiratory syndrome coronavirus 2 (SARS-CoV-2) has emerged in 2019 and rapidly spread worldwide in early 2020^1,2^. Severe prognostic symptoms have become a global problem and the development of effective treatments and drugs is urgently needed^3,4^. Coronaviruses are known for their impact on the respiratory tract, but SARS-CoV-2 can invade and infect the heart, and lead to a diverse spectrum of cardiac manifestations, including inflammation (myocarditis), arrhythmias, heart attack-like symptom, and heart failure^5,6^. The tropism of organs has been studied from autopsy specimens. Sequencing data has revealed that SARS-CoV-2 genomic RNA was highest in the lungs, but the heart, kidney, and liver also showed substantial amounts^7^. In fact, above 1,000 copies of SARS-CoV-2 virus were detected in the heart from 31% patients who died by COVID-19^8^.

The heart is one of the many organs with high expression of SARS-CoV-2 receptor, angiotensin converting enzyme 2 (ACE2)^9^. SARS-CoV use its spike (S) glycoprotein to bind with ACE2, and mediate membrane fusion and virus entry^10,11^. In the process of ACE2 receptor-dependent syncytium formation (i.e., cell-cell fusion), the surface S proteins are cleaved and activated by transmembrane protease serine proteases (TMPRSSs)^11^. The ACE2-dependent SARS-CoV-2 virus entry is reportedly achieved through endocytosis regulated by phosphatidylinositol 3-phosphate 5-kinase (PIKfyve)^12^, the main enzyme synthesizing phosphatidylinositol-3,5-bisphosphate (PI(3,5)P_2_) in early endosome^13^, two-pore channel subtype 2 (TPC2)^12^, a major downstream effector of PI(3,5)P_2_^14^, and cathepsin L^12,15^, a cysteine protease that cleaves S protein to facilitate virus entry in lysosome. In this study, we focused on ACE2 internalization as a therapeutic target of severe COVID-19. Among 13 approved drugs that we have previously identified as potent inhibitor of cardiotoxicity caused by doxorubicin treatment^16,17^, clomipramine, a tricyclic antidepressant^18^, was found to suppress SARS-CoV-2 invasion and infection in TMPRSS2-expressing VeroE6 cells and human iPS-derived cardiomyocytes (iPS-CMs). Clomipramine also synergistically suppresses SARS-CoV-2 virus infection with remdesivir, suggesting the potential as a breakthrough treatment for severe COVID-19.

## Result

### Clomipramine inhibits SARS-CoV-2 pseudovirus-induced ACE2 internalization

Muscular wasting, caused by atrophic remodeling of muscular tissues, is a major symptom of aging (i.e., sarcopenia) and cancer cachexia, and a risk factor for severe COVID-19^19^. We previously identified several approved drugs with potency to inhibit doxorubicin-induced muscular atrophy^20^, and the best potent ibudilast, an anti-inflammatory drug used in the treatment of asthma and stroke, is now under progress of clinical trials for the treatment of acute respiratory distress syndrome (ARDS) in COVID-19 patients^21^. Therefore, we screened another type of drug that has an additional potency to inhibit SARS-CoV-2 spike (S) glycoprotein pseudovirus-stimulated ACE2 internalization. More than 80% of ACE2-EGFP-expressing HEK293T cells showed a marked ACE2 internalization by S protein (50 nM) treatment (Figure 1A). Among top 13 approved drugs, clomipramine, a tricyclic antidepressant that increases plasma serotonin levels^18^, showed the best potency to inhibit the S protein-induced ACE2 internalization (Figure 1B, Table 1). Clomipramine inhibited SARS-CoV-2 pseudovirus infection in a concentration-dependent manner with an IC_50_ of 682 ± 44 nM (Figure 1C). Desmethylclomipramine, a human metabolite of clomipramine, also inhibited the ACE2 internalization with an IC_50_ of 684 ± 46 nM (Figure 1C). HiLyte Fluor 555-labeled S protein was incorporated into ACE2-EGFP-expressing HEK293T cells and inhibited by clomipramine (Figure 1D). Neither the currently proposed SARS-CoV-2 therapeutic agents^21–24^ nor other tricyclic antidepressant agents showed as promising inhibitory effect as clomipramine (Figure 1E, F). We also developed bioluminescence resonance energy transfer (BRET) assay that reflects relative distance between Renilla luciferase–tagged K-Ras (plasma membrane marker) and yellow fluorescent protein (venus)–tagged ACE2 (Figure 1G). The ACE2 internalization caused by pseudovirus infection determined by the decrease in BRET ratio (ΔBRET) was suppressed by clomipramine in a concentration-dependent manner. (Figure 1H). In fact, HiLyte Fluor 555-labeled S protein was incorporated into human induced pluripotent stem cardiomyocytes (iPS-CMs) (Figure 1I), which was significantly suppressed by clomipramine treatment. These results suggest that clomipramine could be repurposed for the treatment with COVID-19 by inhibiting ACE2 internalization caused by SARS-CoV-2 pseudovirus infection.

**Figure 1.**
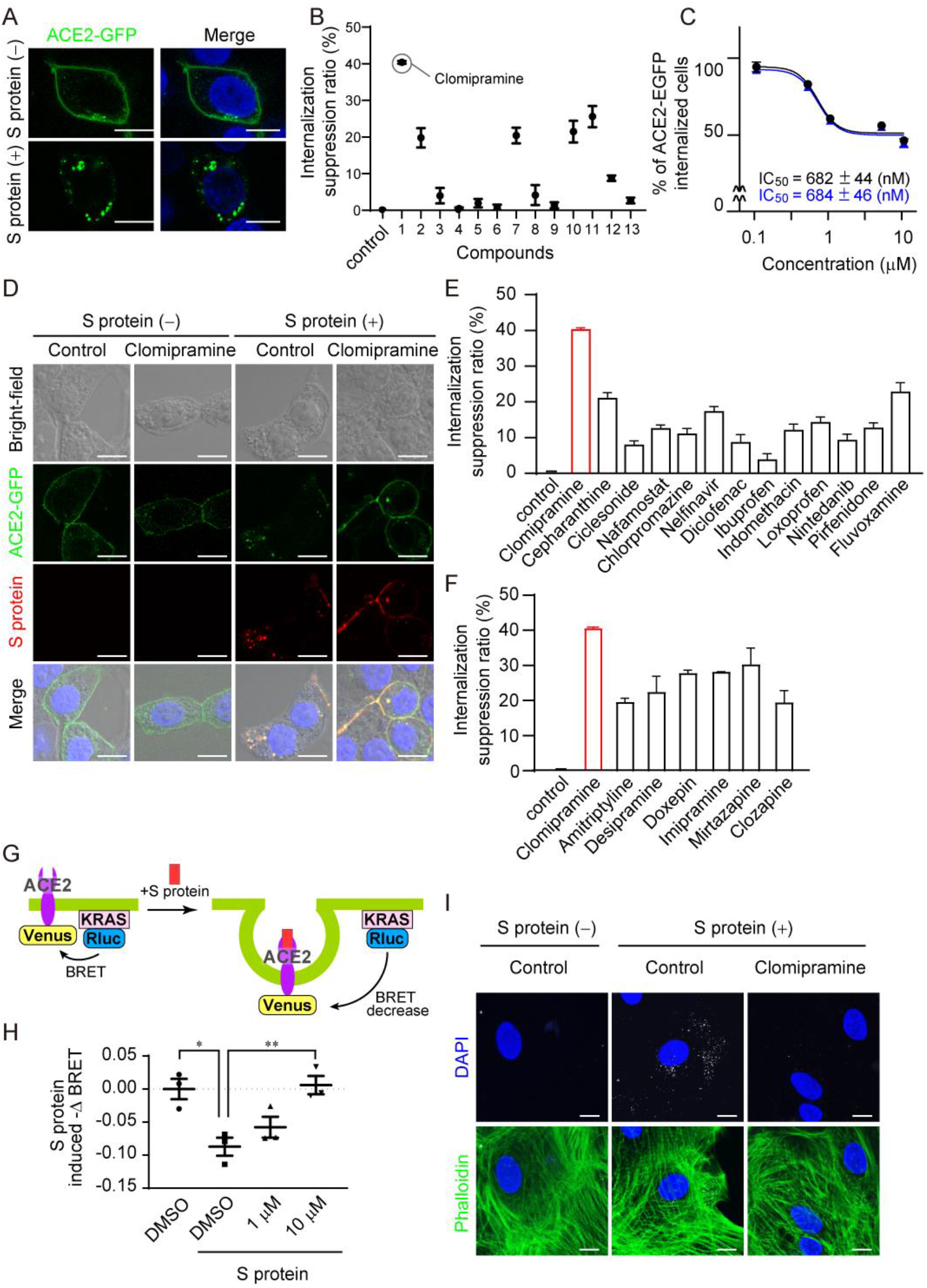
Identification of clomipramine as a potent inhibitor of SARS-CoV-2 pseudo-infection. (A) ACE2 receptor internalization with S protein (50 nM) for 3 hours in HEK293T cells. Scale bars = 10 μm. (B) Inhibitory effects of approved drugs that reportedly have potency to suppress TRPC3-Nox2 complex formation. Cells were treated with each drug (1 μM) 1 hour before exposure to S protein (50 nM) for 3 hours. Compound’s number correspond to Table 1. (C) Concentration-dependent inhibition of ACE2-EGFP internalization by clomipramine and desmethylclomipramine. (D) Internalization of ACE2-EGFP evoked by HiLyte Fluor 555-labeled S protein (50 nM) and its suppression by clomipramine (1 μM) in ACE2-expressing HEK293T cells. Scale bars = 10 μm. (E) Inhibitory effects of repurpose drugs reported to have therapeutic effect targeting SARS-CoV-2. Cells were treated with each drug (1 μM) 1 hour before exposure to S protein (50 nM) for 3 hours. (F) Inhibitory effects of other tricyclic antidepressant agents. Cells were treated with each drug (1 μM) 1 hour before exposure to S protein (50 nM) for 3 hours. (G) The principle of BRET assay. (H) The effect of clomipramine on S protein-induced decrease in BRET ratio (ΔBRET). (I) Protection of human iPS-CMs by clomipramine against pseudo-infection of HiLyte Fluor 555-labeled S protein. Scale bars = 10 μm. All data are shown as mean ± SE; n=3−5 Significance was imparted using one-way ANOVA followed Tukey’s comparison test. **P<0.01, *P<0.05.

**Table 1.** TRPC3-Nox2 complex formation inhibitors^17^.

| Compound No. | Name | Effect | Inhibition of TRPC3-Nox2 complex (%) | Internalization suppression ratio (%) |
| --- | --- | --- | --- | --- |
| 1 | Clomipramine | Inhibitor of reuptake of serotonin and noradrenaline into nerve ending | 54 | 40 |
| 2 | Ibutilast | Phosphodiesterase inhibitor | 75 | 20 |
| 3 | Voriconazole | Inhibitor of synthesis of ergosterol | 66 | 4 |
| 4 | Lynestrenol | Progesterone receptor agonist | 57 | 0 |
| 5 | Pyrilamine | H1 receptor antagonist | 49 | 2 |
| 6 | Clopidogrel | ADP receptor inhibitor | 48 | 1 |
| 7 | Trifluoperazine | D2 receptor antagonist | 47 | 20 |
| 8 | Fipexide | Adenylate cyclase inhibitor | 46 | 4 |
| 9 | Androsterone | positive allosteric effector of GABA<sub>A</sub> receptor | 46 | 1 |
| 10 | Nifedipine | L-type calcium channel blocker | 43 | 21 |
| 11 | Tolnaftate | Inhibitor of synthesis of ergosterol | 42 | 26 |
| 12 | Mifepristone | Progesterone receptor antagonist | 41 | 9 |
| 13 | Ticlopidine | ADP receptor inhibitor | 29 | 3 |

### Clomipramine suppresses SARS-CoV-2 virus infection in human iPS-CMs

Clinically, it is important whether drug treatment after SARS-CoV-2 infection is effective. We confirmed that treatment with clomipramine 1 hour after pseudovirus infection also significantly suppressed the ACE2 internalization (Figure 2A). Therefore, we next examined whether clomipramine prevents SARS-CoV-2 virus infection in TMPRSS2-expressing VeroE6 cells and iPS-CMs. As expected, clomipramine pretreatment prevented the intracellular viral RNA level by 91% (for 6 hours) and 67% (for 18 hours SARS-CoV-2 infection) compared to control in TMPRSS2-expressing VeroE6 cells (Figure 2B-C). In contrast, treatment with ibudilast failed to prevent SARS-CoV-2 virus infection (Figure 2B-D). Treatment with clomipramine simultaneously and 1 hour after SARS-CoV-2 infection also suppressed the intracellular viral RNA level (Figure 2D). Clomipramine showed a concentration-dependent inhibition of SARS-CoV-2 infection with an IC_50_ of 4.4 ± 0.69 μM (Figure 2E). Desmethylclomipramine also suppressed SARS-CoV-2 infection (Figure 2F). Plaque assay revealed that the SARS-CoV-2 virus viability was also reduced by clomipramine, but not ibudilast treatment (Figure 2G). Remdesivir is the RNA-dependent RNA polymerase inhibitor that is approved for the treatment with COVID-19 patients, although World Health Organization (WHO) Solidarity Trial results show little or no effect on hospitalized patients with COVID-19^25^. Therefore, we investigated whether the simultaneous application of clomipramine and remdesivir, which have different targets, would enhance the anti-COVID-19 effect. The single treatment with remdesivir slightly reduced viral RNA level in TMPRSS2-expressing VeroE6 cells, while co-treatment with clomipramine exhibited a dramatic reduction of SARS-CoV-2 viral RNA level (Figure 2H), indicating their synergistic preventing effect against SARS-CoV-2 infection. We further examined whether clomipramine prevents SARS-CoV-2 infection in human iPS-CMs. Indeed, iPS-CMs could incorporate SARS-CoV-2 proteins (Figure 2I). Treatment with clomipramine, but not ibudilast, 1 hour before and after SARS-CoV-2 infection significantly reduced viral RNA levels in iPS-CMs (Figure 2J). These results suggest that clomipramine prevents SARS-CoV-2 infection in TMPRSS2-expressing VeroE6 cells and human iPS-CMs, even after treatment.

**Figure 2.**
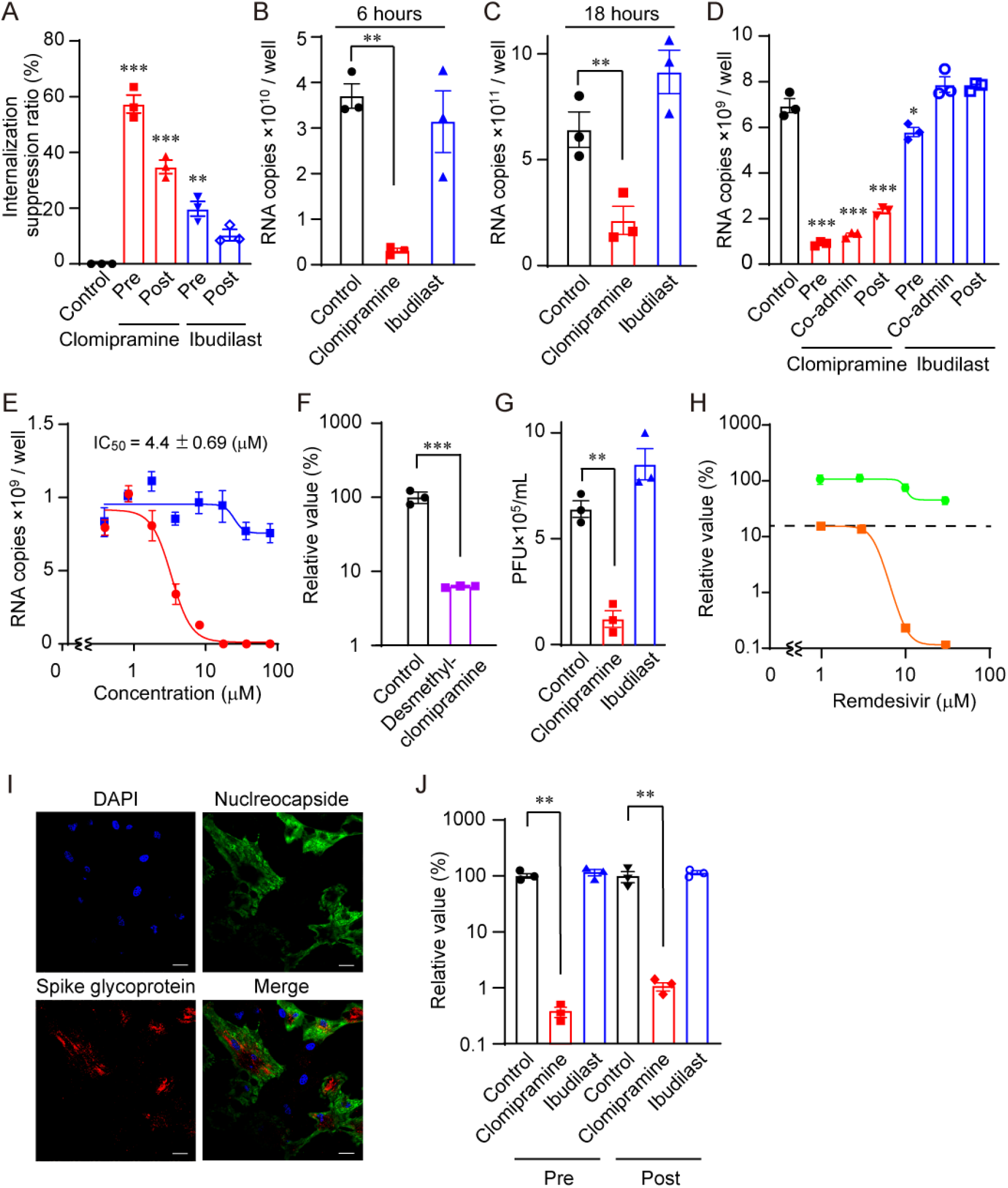
Clomipramine suppresses SARS-CoV-2 infection. (A) Time-dependent inhibition of ACE2-EGFP internalization in HEK293T cells by clomipramine and ibudilast (1 μM) 1hour administered before (Pre) or after (Post) S protein pseudo-infection for 3 hours. (B-C) Effects of clomipramine and ibudilast (10 μM) on SARS-CoV-2 mRNA copy numbers in TMPRSS2-expressing VeroE6 cells exposed to SARS-CoV-2 virus for 6 hours (B) and 18 hours (C). (D-E) Effects of clomipramine and ibudilast (10 μM) on SARS-CoV-2 infection of TMPRSS2-expressing VeroE6 cells. (D) This VeroE6 cells were treated with drugs 0.5 hours before (Pre), simultaneously (Co-admin) or 1 hour after (Post) SARS-CoV-2 infection for 6 hours. (E) Concentration-dependent anti-viral effect by clomipramine (⬤) and ibudilast (◼). (F) Effect of desmethylclomipramine (10 μM) on SARS-CoV-2 infection of TMPRSS2-expressing VeroE6 cells. (G) Results of plaque assay. (H) Combined effect of clomipramine with remdesivir on SARS-CoV-2 infection of TMPRSS2-expressing VeroE6 cells without (⬤) or with (◼) clomipramine (1 μM) 1 hour after SARS-CoV-2 infection. Relative value is SARS-CoV-2 mRNA copy numbers converted with control as 100%. (I) Immunohistochemical images of SARS-CoV-2 incorporation in iPS-CMs. Scale bars = 10 μm. (J) Effects of drugs on SARS-CoV-2 infection of iPS-CMs for 48 hours. All data are shown as mean ± SE; n=3 Significance was imparted using t test for two-group comparisons and one-way ANOVA followed Tukey’s comparison test. ***P<0.001, **P<0.01, *P<0.05.

### Clomipramine non-competitively inhibits the pseudovirus-induced ACE2 internalization

We next investigated how clomipramine inhibits ACE2 internalization caused by pseudovirus infection. The concentration-dependent increase in the number of ACE2 internalized cells due to S protein exposure was suppressed by clomipramine in a concentration-dependent manner, but the mode of inhibition was non-competitive (Figure 3A). Higher concentration (> 10 μM) of clomipramine was required to significantly suppress the interaction between S protein and ACE2, and amitriptyline, another tricyclic antidepressant like clomipramine, never suppress the binding of S protein to ACE2 (Figure 3B). Clomipramine failed to reduce ACE2 enzyme activity (Figure 3C). These results suggest that clomipramine does not seemingly bind with ACE2 but non-competitively inhibits ACE2 internalization in the cells. The receptor binding domain (RBD) of S protein reportedly activates the ACE2-mediated endocytic signal pathway, by which SARS-CoV-2 virus enters the susceptible cells^15^. We first attempted a structure-activity relationship analysis among tricyclic antidepressants with a structure similar to clomipramine, but it was difficult because none of the compounds showed significant inhibition of ACE2 internalization (Figure 3D-F). An *in silico* docking analysis suggests that clomipramine interacts with the S protein at one end of its RBD (Figure 3.G) and that the interacting residues are distinct from other candidate compounds (Figure 3D-F). However, no apparent correlation was observed between the estimated binding energies and the internalization suppression ratio (Figure 3D-E). These results suggest that clomipramine demonstrates a potent anti-COVID-19 effect by non-competitively inhibiting ACE2 internalization induced by S protein pseudovirus.

**Figure 3.**
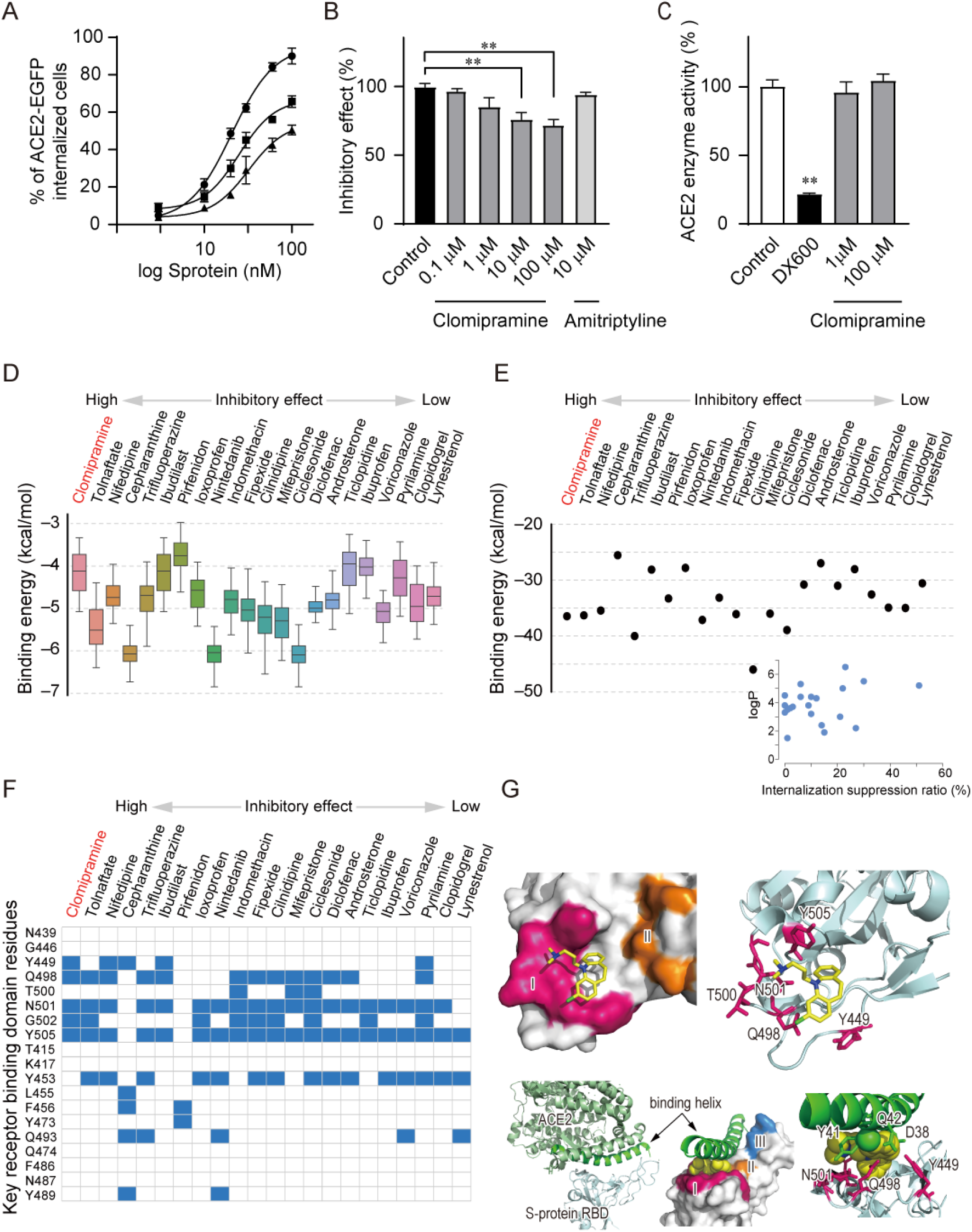
Clomipramine non-competitively inhibits S protein-induced ACE2 receptor internalization. *In-silico* prediction of the binding of 22 candidate compounds to the RBD of S protein. (A) The dose-dependent inhibition of ACE2 receptor internalization by clomipramine (0 μM (⬤), 1 μM, (▲) or 10 μM (◼)) for 1 hour before treatment with S protein (50 nM) in ACE2-EGFP-expressing HEK293T cells. (B) Effects of clomipramine and amitriptyline on the direct interaction between S-protein and ACE2. (C) The effects of clomipramine (1 or 100 μM) and DX600 (1 μM) on ACE2 enzymatic activity. (D) Estimated binding energy distribution of 22 candidate compounds obtained from the docking simulations with SMINA score. The compounds exhibiting more than 20 % of the internalization suppression ratio were listed in red color. No significant correlation with the inhibitory effect is observed. (E) Estimated binding energies of the best poses for 22 candidate compounds minimized using *ad4_scoring*, which includes a desolvation fee energy term. In inset, the logP values of compounds are plotted along the internalization suppression ration (inhibitory effect). The logP and the inhibitory effect weakly correlate and the compounds exhibiting higher inhibitory effect tend to increase the binding energy due to the lesser desolvation penalty. (G) The interaction map of 22 candidate compounds with key residues in the RBD of Spike protein. The residues within 4 Å of any heavy atoms of a drug were colored blue. (H) *In silico* model of the interaction between the RBD Spike protein and clomipramine (colored in yellow). The drug binds to an edge of RBD (region I colored in magenta) (top). The superposition of the docked structure and the crystal structures of ACE2/RBD complex (PDBID: 6m0j) suggests a potential inhibition of ACE2/RBD interaction by the drug (bottom). All data are shown as mean ± SE; n=3 Significance was imparted using one-way ANOVA followed Tukey’s comparison test. **P<0.01.

## Discussion

Small molecule compounds that suppress SARS-CoV-2 virus infection and approved drugs that are expected to repurpose their indications for COVID-19 treatment have been identified one after another, but there is still few effective drug proven in clinical trials established by WHO^21^. Clomipramine, clinically applied as a tricyclic antidepressant, has been used in basic research as a pharmacological tool to inhibit clathrin-dependent endocytosis^26^, and therefore speculated to have antiviral activity^27^. We substantiated that clomipramine suppresses SARS-CoV-2 virus infection and proliferation in human iPS-CMs by preventing ACE2 internalization (Figure 1, 2). In addition to a long half-life pharmacodynamics of clomipramine, its metabolite (desmethylclomipramine) had the similar inhibitory effect of SARS-CoV-2 infection (Figure 1, 2), suggesting that the antiviral effect of clomipramine lasts for a long time. In the process of SARS-CoV-2 virus incorporation into cells with ACE2, serine proteases-dependent plasma membrane pathway (targeted by nafamostat and cepharanthine) and cathepsin-dependent endosome-lysosome fusion pathway (targeted by chloroquine) have been attracted attention as therapeutic targets. In fact, several candidate compounds have also been identified^21,22^. On the other hand, we have not clarified the molecular mechanism underlying suppression of ACE2 internalization by clomipramine. However, while clomipramine demonstrated a potent inhibition of ACE2 internalization due to pseudovirus infection, no drug with the same or better inhibitory effect as clomipramine was obtained (Figure 1). This result may explain why clomipramine potently suppresses SARS-CoV-2 virus infection.

As to the possibility of the improvement of severe COVID-19 by clomipramine, we suggest an anti-inflammatory effect of clomipramine like ibudilast^17^. We have previously reported that transient receptor potential canonical (TRPC) 3 protein, a major component of receptor-activated cation channels, contributes to pathological myocardial atrophy caused by anti-cancer drug treatment^16^, by promoting NADPH oxidase (Nox) 2-dependent reactive oxygen species (ROS) production on cardiac plasma membrane through interacting with Nox2 and preventing Nox2 from ER-associated degradation^20^. Severe COVID-19 symptoms include abnormalities of many organs other than the heart, such as pneumonia, ARDS, olfactory disorder, dysgeusia, vasculitis, headache, and fever^28,29^. Particularly, ibudilast, identified as the best potent inhibitor of TRPC3-Nox2 complex formation, is now under progress of clinical trials for the treatment of ARDS in COVID-19 patients^21^. We have confirmed that clomipramine attenuated the doxorubicin-induced cardiomyocyte atrophy by inhibiting TRPC3-Nox2 complex formation (data not shown). Thus, future studies using COVID-19 model animals will clarify whether clomipramine is effective for the prevention and treatment of severe COVID-19.

Although clomipramine showed a concentration-dependent inhibition of SARS-CoV-2 infection, its mode of action was not competitive inhibition. *In silico* docking analysis predicts that clomipramine interacts with the S protein at the end of the RBD domain, whereby inhibiting ACE2-S protein interaction at higher concentration. However, the binding is weak and therefore this might not fully explain the potent inhibitory effect on SARS-CoV-2 infection by clomipramine. Further study is necessary to identify a molecular target by which clomipramine inhibits ACE2 internalization caused by SARS-CoV-2 virus.

The ACE2 upregulation due to exposure to risk factors for severe COVID-19 is considered as a compensatory mechanism for attenuating the action of angiotensin (Ang) II. Clomipramine does not affect basal ACE2 expression level, but suppresses pathological ACE2 upregulation caused by exposure to risk factors. Additionally, since clomipramine has no impact on ACE2 enzyme activity, it is considered that clomipramine has little effect on AngII and Ang1-7 / Mas receptor signaling pathways in normal cells. On the other hand, it has been reported that Ang1-7 / Mas receptor signaling is attenuated in COVID-19 with ACE2 internalization and degradation^30^. Since clomipramine suppresses ACE2 internalization without reducing ACE2 enzyme activity, clomipramine is also expected to contribute to the maintenance of Ang1-7 / Mas receptor signaling after infection.

In conclusion, this study establishes clomipramine as a potent anti-SARS-CoV-2 agent in human cardiomyocytes. As the co-treatment of clomipramine with remdesivir synergistically suppresses SARS-CoV-2 infection, repurposing of clomipramine will be a promising therapeutic strategy for the prevention and treatment of severe COVID-19.

## Methods

### Cell culture

HEK293 (ATCC, CRL-1573), ACE2-EGFP-expressing HEK293T ^31^ and TMPRSS2-expressing VeroE6 (JCRB, 1819) were cultured in Dulbecco’s modified eagle medium (DMEM) supplemented with 10% FBS and 1% penicillin and streptomycin. Human iPS cardiomyocytes (iPS-CMs) product, iCell Cardiomyocytes^2^, was purchased from FUJIFILM Cellular Dynamics, Inc. and maintained according to the manufacturer’s instructions. Plasmid DNAs were transfected into HEK293 cells with ViaFect Transfection Reagent (Promega) according to manufacturer’s instruction.

### S protein induced ACE2-internalization assay

ACE2-EGFPexpressing HEK293T cells (1.5×10^4^ cells/well) were incubated at least 24 hours before addition of S-protein. In each well, S-protein (50 nM) or HiLyte Fluor 555-labeled S-protein (100 nM) were added and cells were incubated for 3 hours at 37 °C, 5% CO_2_. Cells were fixed in 4% paraformaldehyde for 10 min to stop the reaction and mounted with ProLong Diamond Antifade Mountant containing DAPI (Invitrogen). Imaging was performed on a BZ-X800 microscope (Keyence). ACE2-internalized cells were counted in each section and normalized without S protein as a control.

### S protein & HiLyte Fluor 555-labeled S protein

Recombinant SARS-CoV-2 spike protein (S protein) were purified using the baculovirus-silkworm system^32^. Purified S protein were chemically labeled with a fluorescent probe using HiLyte Fluor 555 Labeling Kit-NH_2_ (Dojindo).

### BRET assay

HEK293 cells were seeded on a 12 well plate, 24 hours before transfection. To analyze the interaction of KRAS and ACE2, BRET between KRAS-Rluc (donor) and ACE2-venus (acceptor) was measured. For the BRET saturation assay, HEK293 cells were transfected with a constant amount of KRAS-Rluc with an increasing amount of ACE2-venus or venus as a negative control. After 24 hours of transfection, cells were suspended in DMEM and were transferred to 96-well plates. In each well, S-protein (50 nM) were added and incubated for 3 hours at 37 °C, 5% CO_2_. Coelenterazine h (5 μM) was added 10 min before BRET measurement. Luminescence and fluorescence signals were detected using Nivo plate reader (PerkinElmer). The BRET ratio was calculated by dividing the fluorescence signal (531/22 emission filter) by the luminescence signal (485/20 emission filters).

### Immunohistochemistry

For immunohistochemistry, iPS-CMs were fixed in 4% paraformaldehyde and then washed twice in PBS. The fixed cells were permeabilized and blocked using 0.1% Triton X-100 and 1% BSA in PBS for 1 hour at room temperature. Alexa Fluor 488 phalloidin (Thermo Fisher Scientific) was diluted with PBS and reacted for 1 hour at room temperature. After mounting with ProLong Diamond Antifade Mountant containing DAPI (Invitrogen), images were captured using a confocal laser-scanning microscope (LSM700, Zeiss, Germany).

### SARS-CoV-2 infection

SARS-CoV-2 strain JPN/TY/WK-521 was distributed by National Institute of Infectious Diseases in Japan. TMPRSS2 expressed VeroE6 cells (1.5×10^4^ cells/well) were seeded in 96-well plates 24 hours before infection. Each compound (10 μM) was added. SARS-CoV-2 virus were added at multiplicity-of-infections (MOI) 0.05 or 0.1. iPS-CMs (3×10^4^ cells/well) were seeded in 96-well plates and were cultured for 5-7 days before infection. Each compound (10 μM) was added and SARS-CoV-2 virus were added at MOI 2.5. After infection, intracellular RNA was extracted with CellAmp direct RNA prep kit (TAKARA) according to the manufacturer’s instructions. Quantitative real-time PCR was performed using a TaqMan Fast Virus 1-Step Master Mix (Thermo Fisher Scientific), 2019-nCoV RUO Kit (Integrated DNA Technologies) and 2019-nCoV_N positive Control (Integrated DNA Technologies) with a QuantStudio 7 Flex Real-Time PCR System (Thermo Fisher Scientific).

### Plaque assay

Plaque assay was performed essentially as described^33^.Infection medium of TMPRSS2-expressing VeroE6 cells after SARS-CoV-2 virus infection was diluted by DMEM supplemented with 2% FBS and 1% penicillin and streptomycin. This sample spread on an agar plate with 2% FBS and 1% methylcellulose and cultured at 37 °C for 3 days. VeroE6 cells on the plate were fixed in 10% neutral-buffered formalin (Fujifilm Wako Chemicals, Tokyo, Japan), and strained with methylene blue. Numbers of the plaque in each well were counted to calculate infective per one ml of the samples.

### Immunohistochemistry – SARS-CoV-2

Human iPS-CMs (3×10^4^ cells/well) were seeded. SARS-CoV-2 were added at MOI 2.5. Forty-eight hours after infection, cells were fixed in 4% paraformaldehyde. The fixed cells were incubated with anti-SARS-CoV-2 nucleocapside (1:100; GeneTex), anti-SARS spike glycoprotein (1:100; abcam) and DAPI. Images were captured using a Nikon A1 confocal microscope (Nikon).

### ACE2 / SARS-CoV-2 Spike Inhibitor Screening Assay

The ACE2:SARS-CoV-2 Spike Inhibitor Screening Assay Kit (BPS Bioscience #79936) was purchased and measured the binding of ACE2 and SARS-CoV-2 Spike in the presence and absence of inhibitors, according to the manufacturer’s instructions.

### ACE2 inhibitor Screening Assay

The ACE2 Inhibitor screening assay kit (BPS Bioscience #79923) was purchased and measured the exopeptidase activity of ACE2 in the presence and absence of inhibitors, following the manufacturer’s instructions.

### *In silico* binding mode prediction of ACE2 / SARS-CoV-2 Spike Inhibitors

The bindings of 22 candidate compounds, including clomipramine, toward the receptor binding domain (RBD) of SARS-CoV-2 Spike protein were analyzed using SMINA program with its own score function (SMINA score)^34^. The structure of the receptor (RBD of Spike protein) was taken from the crystal structure (PDBID: 6m0j)^35^ and hydrogens were added by *h_add* function in PyMOL^36^. SMILE stings of 22 compounds from PubChem^37^ were converted to three-dimensional structures using OpenBabel^38^, where some of the compounds were either protonated (clomipramine, trifluoperazine, pyrilamine, nintedanib, sefarantin and ticlopidine) or de-protonated (dicrofenac, ibuprofen, indomethacin and loxoprofen) using RDKit^39^ according to the pka calculations with ChemAxon software MARVIN^40^. Each compound was initially docked into the region centered at Gln493 of the RBD (30 Å ×30 Å×30 Å), and we used the results as regions for subsequent docking. The procedure was repeated three times to sample 183 bound poses distributed over the RBD surface interfacing ACE2 / SARS-CoV-2 binding. The estimated binding energy distribution of 22 compounds was shown in Supplemental Figure 5A and the binding mode with the lowest binding energy for clomipramine was shown in Figure 4D. The estimated binding energies of the best poses for 22 compounds were further refined by minimizing using *ad4_scoring*, which corresponds to the AutoDock 4 score including a desolvation fee energy term (Supplemental Figure 5B).

## Materials

Clomipramine, voriconazole, clopidogrel, trifluoperazine, androsterone, nifedipine, ibuprofen, loxoprofen and fluvoxamine were purchased from Fujifilm Wako Chemicals, (Tokyo, Japan). Ibudilast, lynestrenol, pyrilamine, tolnaftate, mifepristone, ticlopidine, nafamostat, chlorpromazine, nelfinavir, diclofenac, indomethacin, nintedanib and pirfenidone were purchased from Tokyo Chemical Industry (Tokyo, Japan). Fipexide, ciclesonide and desmethylclomipramine were purchased from sigma (St.louis, MO, USA). Cepharanthine was purchase from Cayman chemical (Michigan, USA).

### Statistics

G*Power3.1.9.2 software was used to calculate the sample size for each group. All results are presented as the mean ± SEM from at least 3 independent experiments and were considered significant if P < 0.05. Statistical comparisons were made using unpaired t test for two-group comparisons or using one-way ANOVA with Tukey’s post hoc test for comparisons among 3 or more groups when F achieved was P < 0.05, and there was no significant variance in homogeneity. Statistical analysis was performed using GraphPad Prism 8.0 (GraphPad Software, LaJolla, CA). Some results were normalized to control to avoid unwanted sources of variation.

## Acknowledgments

We appreciate the technical assistance from The Research Support Center, Research Center for Human Disease Modeling, Kyushu University Graduate School of Medical Sciences. This work is supported by the grants from Smoking Research Foundation (to M.N. and Ya.K.), “COVID-19 Drug and Vaccine Development Donation Account” Project from Sumitomo Mitsui Trust Bank (to M.N. and T.K.), Uehara memorial foundation (to Yu.K.), Foundation of Kinoshita Memorial Enterprise and Suzuken Memorial Foundation (to K.N.) and in part by JST, CREST Grant Number JPMJCR2024 (20348438 to M.N.), Platform Project for Supporting Drug Discovery and Life Science Research (BINDS) from Japan Agency for Medical Research and Development (AMED) under Grant Number JP20am0101091 (to M.N.) and AMED under Grant Numbers JP18mk0104117 and JP20fk0108263 (to Ya.K.).

## Author contributions

Yu.K., K.N., S.R., K.M. and M.N. wrote the paper, Ya.K. and Yu. K. designed experiments, Yu.K., S.Y., K.N., A.S., D.T., J.L., K.Y. and H.A. performed experiments, S.R. and K.M. performed *in silico* analysis, T.K., N.K., Y.Ib. and Y.Im. contributed new reagents/analytic tools and critical suggestions, T.T., S.R., K.M. and A.N. analyzed and interpreted data, Ya.K. and M.N. edited the paper.

## Notes

### Competing Interest Statement

The authors have declared no competing interest.

